# Multi-variate statistical and machine learning reveals the interplay between sex and age in antibody responses to *de novo* SARS-CoV-2 infection and vaccination

**DOI:** 10.1101/2023.12.05.569965

**Authors:** Miroslava Cuperlovic-Culf, Steffany Bennett, Yannick Galipeau, Pauline S. McCluskie, Corey Arnold, Salman Bagheri, Curtis L. Cooper, Marc-André Langlois, Jörg Fritz, Ciriaco A. Piccirillo, Angela M. Crawley

**Author notes:** **co-senior authors** (Miroslava Cuperlovic-Culf, Jörg Fritz, Ciriaco A. Piccirillo, Angela M. Crawley).

## Abstract

Prevention of negative COVID19 infection outcomes and infection/vaccine-acquired immunity is associated with the quality of antibody responses, whose variance by age and sex are poorly understood. Integrated, network approaches, identified sex and age effects in antibody responses and neutralization potential of *de novo* infection and vaccination throughout the Covid-19 pandemic. Cluster analysis found neutralization values followed SARS-CoV-2 specific receptor binding RIgG, spike SIgG and S and RIgA levels based on COVID19 status. Stochastic behavior tests and other analytical methods revealed sex differences only in persons <40y.o. Serum IgA antibody titers correlated with neutralization only in females 40-60y.o. Network analysis found males could improve IgA responses after vaccination dose 2, unlike >60y.o. females. Complex correlation analyses found vaccination induced less antibody isotype switching and neutralization in older persons, especially in females. Sex dependent antibody & neutralization behavior decayed fastest in older males and with vaccination. Such sex and age characterization by machine learning can direct studies integrating cell mediated responses to define yet elusive correlates of protection and inform age and sex precision-focused vaccine design.

## Introduction

Sex and age differences in anti-viral responses, vaccination, and long-term recovery are increasingly recognized in COVID19 ^1,2^ as key determinants to disease susceptibility and severity. During viral infections, females have greater inflammatory, antiviral, and humoral immune responses compared to males. Early in the COVID19 pandemic it was observed that infection prevalence is sex-independent or possibly slightly higher in females. In contrast, males are more likely to experience severe disease, combined with increased hospitalization rates and higher mortality^3,4^. Increased disease severity was particularly significant in men over 60 years of age^2^. In a large, hospitalized patient cohort study of 308,010 patients, the mortality rate was higher in males regardless of age, race, or comorbidities^5^. This is in agreement with other coronaviruses including the Middle East respiratory syndrome (MERS) epidemic, where mortality rate is also higher in males^6^. Differences in sex-related mortality risk have been observed in animal models of acute respiratory viral infections^7^, further suggesting significant sex effects influencing the outcomes. The molecular basis for these divergent immunological outcomes in males and females is thought to be related at least in part to differences in sex steroid synthesis, which can have a crucial role on the function and regulation of the immune response^8,9^. Indeed, sex steroids have been reported to exert a suppressive role in immune functions suppressing immune cell activity by promoting anti-inflammatory mediators. However, the nature and dynamics of antibody production throughout infection and vaccination in men and women is not well understood^10^.

Adaptive immunity to SARS-CoV-2 infection depends on the concerted action of both humoral and cell-mediated immune components. IgG, IgM, and IgA against spike and RBD are detected in plasma 7-8 days following the onset of symptoms in infection with IgA persisting for over 400 days after symptom onset^11^. Detectable IgM and IgG Abs occur concurrently with increased plasma B cells. While IgG levels reportedly decline modestly ∼8 weeks after symptom onset, recovered patients maintain high spike-specific IgG and IgA titers. Neutralizing Abs (nAbs) as well as binding Abs (bAbs) can control SARS-CoV-2 infection; convalescent plasma containing nAb and bAb administered as a therapeutic can improve clinical symptoms^12^. In SARS-CoV-2 convalescent subjects, IgG1, IgA1 and IgM antibodies against spike and RBD all have capacity for virus neutralization^11^. Based on regression analysis, IgM and IgG1, show the strongest contribution to neutralization, however direct analysis shows that IgA was also able to neutralize^11^.

Severe COVID19 patients display an acute respiratory distress syndrome (ARDS), which can be mediated directly by the virus and/or a hyper-active immune response. SARS-CoV-2 antigen-specific CD4^+^ and CD8^+^ T cells have been identified in recovered and active disease patients and contribute to protection from re-infection^13–16^. The immune profiles of COVID19 patients with moderate disease indicate a protective T cell response, while patients with severe disease exhibit exaggerated systemic inflammation, signs of T cell exhaustion and lymphopenia, and decreased polyfunctionality or cytotoxicity, all features of dysregulated adaptive immunity. Different immunotypes predict the course of infection, whereby coordinated action between CD4^+^, CD8^+^ T cells, and Ab contribute to optimal viral control, efficient immune protection, and better disease prognosis.

Viral-induced inflammation increases with age and is associated with a hyper-inflammatory response leading to significant lung damage as well as increased morbidity and mortality rates in older adults. Elevated levels of inflammatory cytokines (e.g., IL-6), increased inflamm-aging and T cell senescence, as well as higher titers of anti-S/-N IgG and IgM correlate with worse clinical readouts with older age, suggesting adverse effects of the immune response on disease severity^14,17,18^.

Immune responses are known to differ in males and females leading to sex-specific responses to vaccination and infections^19^. Female mice have a more robust humoral and cellular immunity than their male counterparts following a primary challenge with influenza virus whereas vaccination yields equivalent protection in both sexes^20^. Transfer of immune sera from vaccinated females to naïve mice provided better protection compared to transfer of sera from vaccinated males. This superior sex-associated protection is thought to be a consequence of an increase in antibody titers, IgG antibodies of greater neutralization capacity, and effective immunoglobulin (Ig) isotype switching in females contributing to more effective antibody function and neutralization capacity. A meta-analysis included all peer-reviewed articles showing serum levels of IgA, IgG, or IgM in adult human beings^21^ (117 articles) suggests that older age and male sex are associated with higher vaccine-specific IgA levels and that lower IgM and IgG levels were not influenced by age and sex. A population-based study^22^ support these findings. Increased levels of serum IgA and decreased IgM with age and higher IgA titers in males than females were noted. Interestingly, IgG level decrease with age (up to 60)^22^.

In an effort to better understand the observed sex and age differences in SARS-CoV-2 infection risk, disease outcome, and response to COVID19 vaccination, we applied a novel network analysis strategy^23–25^ to track the evolution of antibody generation and functionality following natural infection and/or vaccination in a large longitudinal cohort referred to as *Stop the Spread Ottawa* (SSO)^26^, pinpointing changes in the response to infection and/or vaccination in immune-competent or immune-compromised individuals.

## Materials and Methods

### Serology assay

Antibodies against SARS-CoV-2 were measured using a high-throughput direct chemiluminescent ELISA performed on MicroLab STAR robotic liquid handlers (Hamilton) fitted with a 405TS/LS LHC2 plate washer (Biotek/Agilent) (full methods described previously)^27^. Briefly, SARS-CoV-2 antigens (full trimeric Spike (Wuhan), receptor binding domain (RBD) (Wuhan) and the Nucleocapsid protein) were generously provided by Dr. Yves Durocher (National Research Council of Canada (NRC), Montréal) or purchased from the Metrology division of NRC and coated on assay plates. Human serum samples or controls were either diluted 1:100 for measurement of all antibody-isotype combinations, or 1:10,000 to resolve saturating signals for Spike and RBD due to vaccination. Controls included an isotype-antigen specific calibration curve pooled human sera from SARS-CoV-2 naïve, convalescent, vaccinated individuals as well as a general immunoglobulin control. Isotype-specific secondary antibodies detected bound Igs. Antibody titers were determined as lab-specific ug/mL concentrations in relation to the isotype-antigen specific calibration curve or, for IgG, converted to international units (binding antibody units (BAU)/mL) via modelized WHO Standard (NIBSC 20/136).

### Neutralization assay

A surrogate protein-based neutralization assay was used to determine the neutralization efficiency in sera. The detailed methods, validation and calibration of the assay is available here^27^. Briefly, SARS-CoV-2 full-trimeric Spike (Wuhan)-coated plates were blocked, and then human serum samples and controls were assayed. Controls included a calibration curve (NRCoV2-20-Fc, NRC) as well as pooled human sera from naïve, convalescent, and vaccinated individuals. Subsequent incubation with biotinylated ACE2 (NRC) followed by streptavidin-HRP linkage and chemiluminescent substrate addition permitted the determination of a luminescent signal. Raw luminescence values were blank (serum-free, no ACE2) adjusted using ACE2-only wells (representing unimpeded ACE2-Spike interaction). Percentage inhibition was established by measuring the reduction in ACE2-Spike interaction (0% inhibition = maximal ACE2-Spike interaction)

### Samples

All biological specimens from SSO are stored in the Coronavirus Variants Rapid Response Network (CoVaRR-Net) Biobank, following the highest standards of biobanking practices in sample collection, processing, storage, access, and distribution (upon application). Sample collection followed approved ethics protocols (REB # H-09-20-6135).

### Knowledge discovery in data

#### Data preprocessing

Data points generated from serology and neutralization assays that were below the level of detection were imputed using 1/5 of the lowest measurement for the given feature. Values above the upper level of quantification were imputed with the highest measured value for the assay. In some analyses, data for each feature across samples is normalized or log transformed, and this transformation is indicated. No other data manipulation was performed.

#### Data Overview

General overview of the serology measurements was performed using Principal Component Analysis (PCA), guided PCA (gPCA), t-distributed stochastic network embedding (t-SNE) and hierarchical cluster (HCL) analysis. Cluster analysis was done using Fuzzy C-means clustering. Briefly this approach determines the level of belonging of each feature to each cluster in this way allowing for possible multiple groupings. Fuzzy c-means (FCM) and related methods fuzzy j-means^28^ allows each feature to belong to more than one group by providing a degree of membership, “belonging”, to each cluster by maximizing proximity between similar features and distance between dissimilar features. FCM is based on the minimization of the objective function: 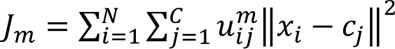 where *m* ∈ (1, ∞) is the “fuzzyfication” factor, 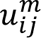 is the membership degree for feature *x*_*i*_ to the cluster *j* with *c*_*j*_ defining the cluster center. FCM clustering assigns objects to groups with features belonging to the same clusters showing more similarity to each other than to features in other clusters. Higher membership value indicates stronger belonging to the cluster with membership value of 1 ultimately indicating that feature is only associated with the single cluster. An optimal number of clusters in FCM analysis was determined using within-cluster sum of squares and silhouette methods applied to k-means clustering as a directly related crisp method. Clustering with this approach was performed to determine grouping of features in different subject cohorts.

Guided PCA (gPCA) was performed following method previously presented for identification and batch analysis^29^. Briefly, this approach is focusing on the investigation of the importance of each group corresponding to the corresponding principal component (PC). In order to determine group loading contribution “batch” in our case group indicator matrix G, defined as *n* x *b* matrix where *n* is number of samples and *b* number of groups as assembled such as: *g_ik_* = 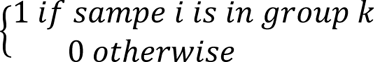. Then single value decomposition of the guided matrix calculated as *n*′*b* gives group loadings in PCA and can be used to determine separation of groups in a guided sense.

#### Correlation analysis

Correlations were calculated using Pearson, distance and point bi-serial correlation. Pearson correlation is a standard method used for linear correlation assessment and was used for the determination of correlation between distance correlation level values for different subgroups of the cohort.

**Distance correlation** was performed using previously published applications^25^. Briefly, distance correlation^30^ was calculated using distance covariance, rather than direct covariance used in Pearson method. In this way, distance correlation provides information about both linear and non-linear correlations. Formally, distance correlation between features *X* and *Y* is calculated as:

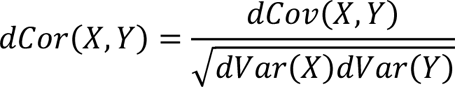

distance covariance is determined as:

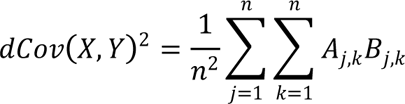

where *A* and *B* are calculated as simple linear functions of the pairwise distances between elements in samples *X* and *Y*. *A* and *B* are doubly centered distance matrices for variables *X* and *Y,* respectively, calculated from the pairwise distance between elements in each sample set calculated as:

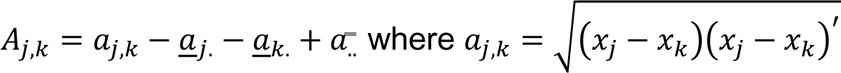

and <u>*a*</u>_*j*._ and <u>*a*</u>_*k*._ are respectively the *j*-row and *k*-column mean values and *a̿*_.._ is the overall mean of matrix *A*. The distance between *x*_*j*_ and *x*_*k*_ is calculated using Euclidian distance. Matrix *B* is populated using equivalent measures for variable *Y*.

**Square distance correlation**, i.e., the coefficient of determination, is used to represent the fraction of the variation in a variable that can be explained by the other variable. Relative square distance correlation is calculated as a square distance correlation divided by the sum of square distance correlation values for all features.

Correlation between binary values (e.g., sex) and continuous values (e.g., serological measurements) was determined using **Point-biserial correlation** method as:

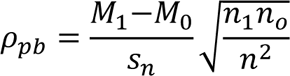

where correlation is calculated between binary and continuous variable by dividing continuous variable into two groups, 0 and 1, based on values of the binary feature and where *M0* and *M1* are mean values, *n0* and *n1* are numbers of samples of two subgroups and *sn* is the standard deviation for the complete continuous variable set.

**Polynomial function** regression analysis was performed using second order polynomial and exponential functions and bisquare robustness analysis. Briefly, the bisquare robustness method for polynomial fit determination minimizes a weight sum of squares with points near the line getting full weight and points further from the line getting a reduced weight. Points that are farther from the line are expected by random chance and hence are given a weight of zero. In this way, the bisquare method aims to find a function of best fit using a least squares approach while at the same time minimizing the effect of outliers.

#### Feature selection

Selection of the most significantly different features between different groups of samples was performed using three different methods to determine method-independent major features: 1) statistical feature selection using F-test (function *fsrftest* running under Matlab); 2) machine learning methods Relieff (function relief in Matlab); and 3) neighborhood component analysis for regression (*fsrnca* function in Matlab). Features were considered significantly different if selected by all three methods. Data was z-score normalized for each feature prior to analysis. In F-test features with p<0.05, −log(p)>3 is selected as significant. In Relieff, all features with positive weight are determined as possibly contributing to group selection. Similarly, in fsrnca, all features with positive weights are kept as possibly significant. *Fsrnca* aims to find, i.e., learn, a distance metric to maximize leave-one-out classification performance. Thus, the feature selection analysis used here combines two different ML methods including filter (Relieff) and wrapper (fsrnca) with a statistical method (F-test) to determine consistently significant features.

## Results

### Age and Sex weakly affect overall response to vaccination and natural infection

In this analysis, serially-collected serum samples from 970 participants (3666 samples in total) were separated into group by the vaccination period, age, and sex of the participant as well as by SARS-CoV-2 infection-acquired immunity status (Table 1). Nine different antibody responses were measured from each serum sample corresponding to IgG, IgA, and IgM titers specific for SARS-CoV-2 receptor binding domain (R), spike (S) and nucleoprotein (N) as well as neutralization measurements.

**Table 1.** Overview of study groups and time points assembled for this data analysis.

|  | **Unvaccinated**<br>**(male/female)** | **Dose 1**<br>**(male/female)** | **Dose 2**<br>**(male/female)** |
| --- | --- | --- | --- |
| **Unique study participants (n=970)** | 198 / 394 | 122 / 244 | 140 / 290 |
| **Samples** | 393 / 643 | 205 / 422 | 440 / 1012 |
| Positive | 177 / 163 | 84 / 90 | 119 / 168 |
| Negative | 216 / 480 | 121 / 332 | 321 / 844 |
| <40 y.o. | 99 (30.3±4.5) / 249<br>(30.6±4.9) | 44(30.4±5.2) / 118<br>(31.2±4.8) | 111(31.5±4.6) / 368<br>(31.4±4.5) |
| Age [40-60) y.o. | 173(48.0±6.3) /<br>275(49.1±6.2) | 53(49.6±5.7)/<br>134(50.6±5.60) | 172(46.8±5.4) /<br>411(50.3±6.0) |
| >=60 y.o. | 121(66.6±4.8) /<br>119(66.0±4.3) | 55(68.0±5.0) /<br>67(65.7±4.7) | 104(65.7±5.2) /<br>135(65.8±4.7) |

Dose #: 0 unvaccinated; Dose 1 and 2: minimum day 14 post-vaccination and before next dose Positive: seropositive for IgG-N (nucleoprotein); Negative: seronegative for IgG-N; y.o.: years old

An initial data overview was performed for the complete set using unsupervised visualization and clustering methods to determine dominant patterns in the data. Principal Component Analysis (PCA) (Supplementary Figure 1A) indicates a dominant effect of vaccination on serology profiles in all age groups with the most apparent separation between Unvaccinated and Vaccination 1 groups. Change over time with vaccination is once again significantly different between COVID19+ and COVID19-groups, showing major value difference in SIgG between these two groups as well as lower level of neutralization in COVID19-cohorts in all age groups and both sexes even after vaccination (Figure 1A). However, from observing individual serology factors difference between age and sex groups in not apparent. However, effect of age and sex on the serology measure starts to be visible in the ANOVA analysis (Figure 1B) as well as guided PCA (gPCA) analysis, semi-supervised method that shows group loadings in the PCA (Figure 1C) (Reese, Archer et al. 2013). Furthermore, t-SNE analysis (Supplementary Figure 1B), providing projection of sample points while preserving relationship between sample points indicates some grouping of sample points based on sex as is shown with p-values of the significance of the separation by sex in different subgroups (Supplementary Figure 1C). The association of the effect of sex on antibody responses was particularly significant between male and female participants <40 y.o. who were either SARS-CoV-2 antibody negative or positive across all three first doses of vaccination. Meanwhile, the 40-60 y.o. group showed significance only in the SARS-CoV-2 positive subjects after vaccination 1 and 3 and there were limited/transient sex effects in the >60 y.o. group, reserved for SARS-CoV-2 seropositive individuals after the second vaccination dose. Thus, t-SNE suggests that age is a primary determinant of antibody response with complex dependence between the response and sex.

**Figure 1.**
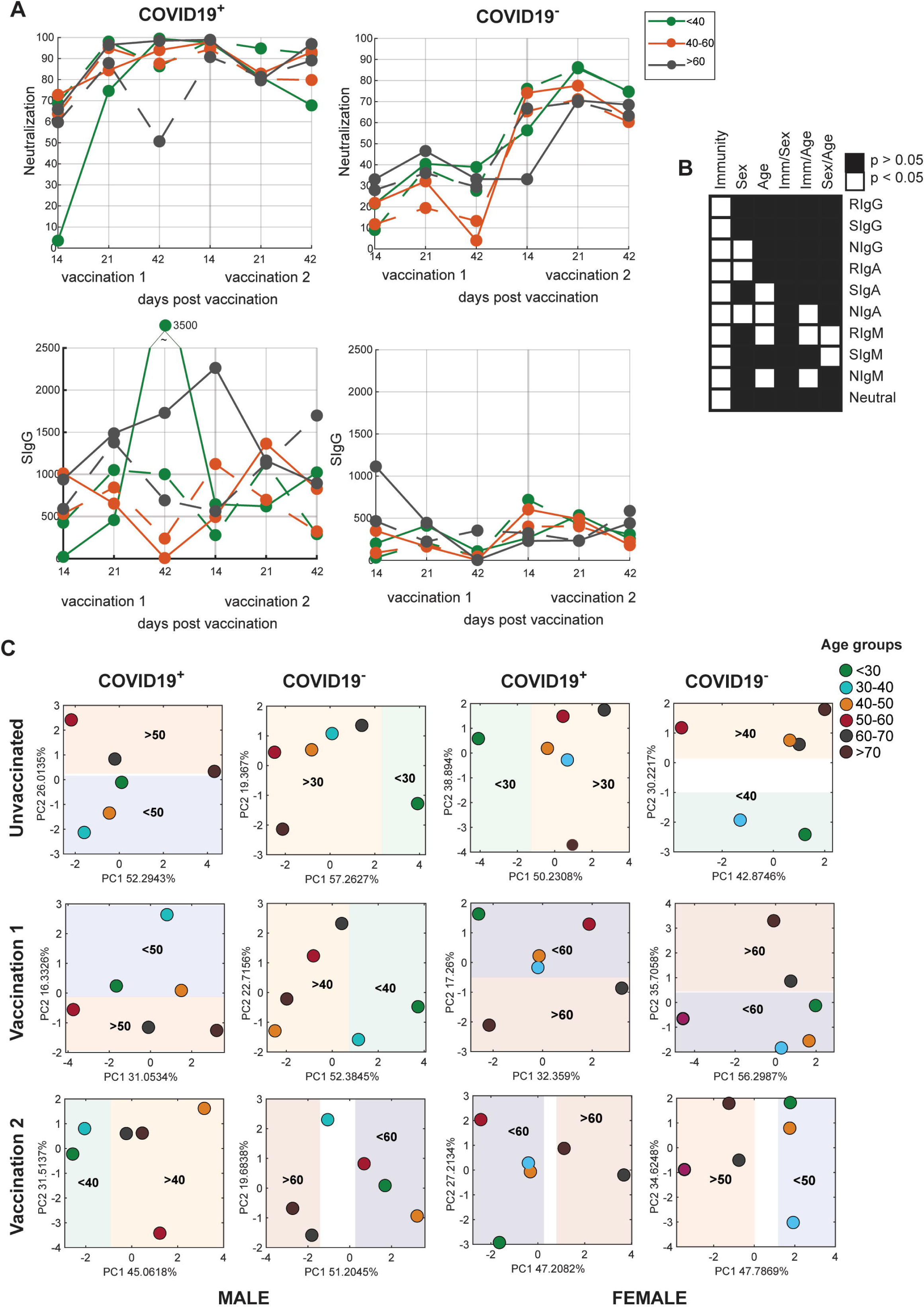
Small differences between sex and age groups in response to vaccination and infection are observed. Samples are divided by COVID19 status (COVID19^-^ = no COVID19 infection history, confirmed by negative NIgG serology; COVID19^+^ = history of COVID19 infection, confirmed by positive NIgG serology). **A)** graphical representation of the mean values for Neutralization and SIgG at three time points after vaccination 1 and vaccination 2 (days 14, 21, and 42 post-vaccination) in cohorts grouped by sex and age (in three age groups <40, 40-60, and over 60 years of age) **B)** 3-way ANOVA evaluation of the mean differences between measurements grouped by acquired immunity (COVID19^-^ or COVID19^+^), sex (male/female) or age (under 40, 40-60, or over 60 years of age). Indicated as white boxes are statistically significant mean differences for direct and combined effect. **C)** gPCA for log transformed and z-score scaled dataset of 9 different antibody responses and Neutralization (10 features) with included PC1 for guided PCA (gPCA) for age groups for each subset divided by vaccination dose and sex; age groups are under 40, 40-60, and over 60 years of age. Shown are loadings for each group.

Further, Hierarchical Cluster Analysis, (HCL, Supp. Figure 2) indicates comparable separation between samples collected pre-and post-vaccination. In all four groups of samples (including male and female groups distinguished by SARS-CoV-2 infection acquired immunity status), HCL shows that neutralization values cluster most closely with RIgG, SIgG, and SIgA levels. In COVID19 positive samples, neutralization is also clustered with RIgA. Uniquely in HCL of COVID19 negative males, NIgG clusters with NIgM, SIgM and RIgM while it is most strongly grouped with NIgA, RIgA, and SIgA in the other groups (Figure 1B). Once again, these clustering results suggest a level of sex dependent behaviour in the relationships between measured antibody responses (Supplementary Figure 2).

### Co-behaviour between antibody measures and neutralization is sex-, age- and immunity-dependent

Detailed analysis of feature co-clustering, i.e., similarities in behaviour between neutralization and antibodies was performed using the Fuzzy C-Means (FCM) clustering method. This approach provides information about the level of co-clustering for each feature to each group, allowing for possible multiple relationships. Data in Figure 2 shows a weight for each antibody response feature for belonging to the cluster relative to their respective neutralization value. Cluster analysis is performed separately for subject groups divided by age, sex, and COVID19infection-acquired immunity status. Unlike HCL analyses of all age groups combined (Figure 1), this approach is applied to three age groups separately and shows differences between males and females, particularly in older and COVID19 positive groups. Interestingly, in females >40 years old RIgA and SIgA co-cluster with neutralization, and males co-cluster with RIgM and SIgM responses. In the <40y.o. groups, SIgG and RIgG levels correlate with neutralization in both males and females.

**Figure 2.**
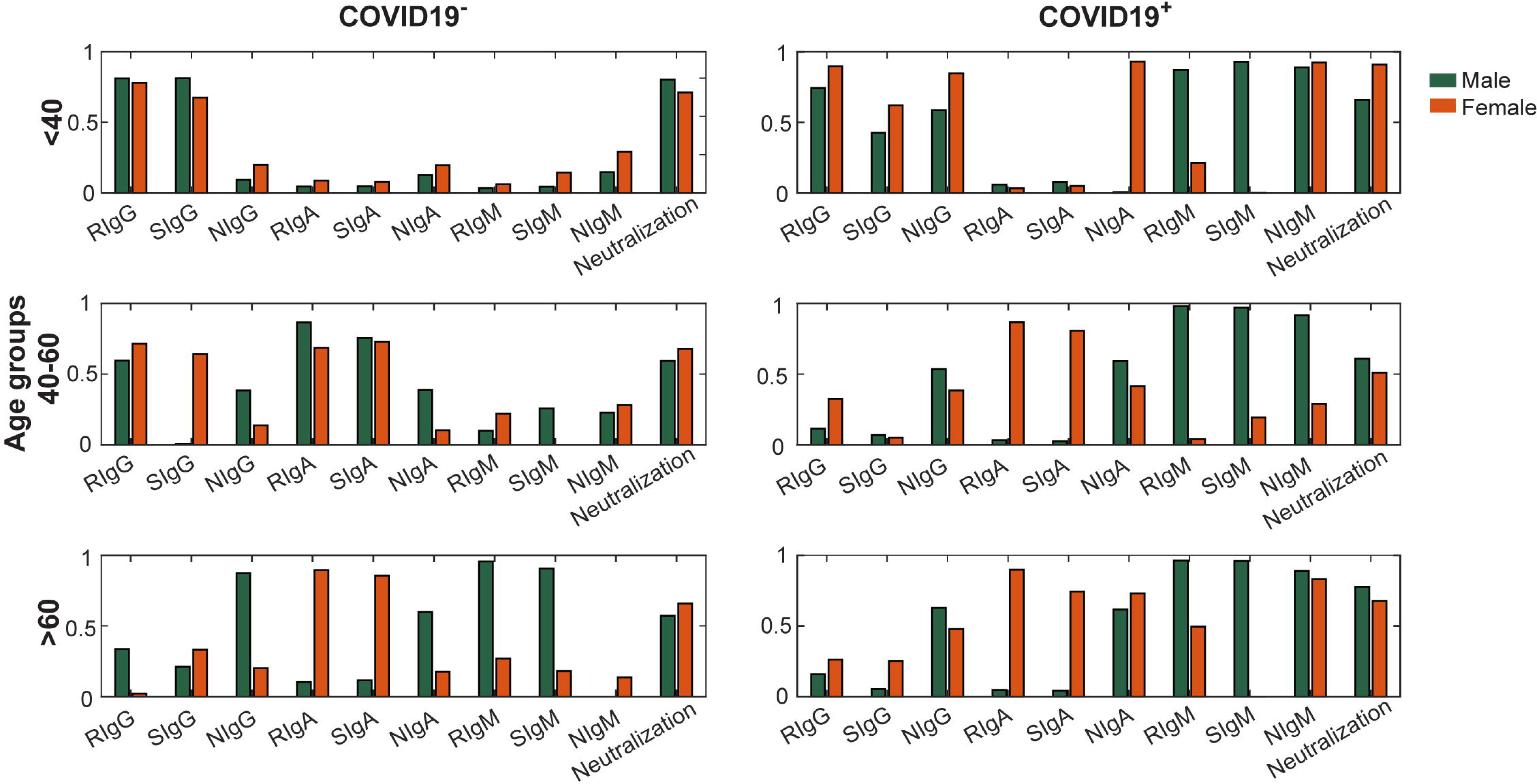
Cluster analysis of antibody responses relative to neutralization shows differences in co-behaviour in males and females by age. Fuzzy C-means clustering of antibody response features across data in subgroups divided by age, sex (male in black, female in orange), and COVID19 infection status (negative or positive SARS-CoV-2 history, confirmed by NIgG level). Shown are membership values, i.e., cluster association of features in different subject cohorts with clusters that have the strongest neutralization membership. Fuzzy C-means here clusters features separately for each of the groups of data divided by age, sex, and COVID19 infection status. Shown are weights (0.0-1.0), i.e., cluster belongings for each antibody response feature relative to neutralization values (weights approaching 1.0 indicate strong clustering with neutralization).

### Major differences in antibody responses between males and females are dependent on age and infection-acquired immunity

We used three different feature selection methods (see Material and Methods) to determine the most significant feature differences (Figure 3). Features that are selected by all three methods are indicated. The data for all time points were combined in the analysis but a separate analysis was performed for COVID19 negative and positive samples across the age groups and is reported here (Supplementary Figures show feature selection in each vaccination group). Differences in IgG and IgA factors are identified in participants <40y.o. COVID19 negative group. Differences in IgM levels are noted in individuals 40-60 y.o. and >60 y.o. The features selected as statistically different using these methods are largely in agreement with features showing major clustering differences in Figure 2.

**Figure 3.**
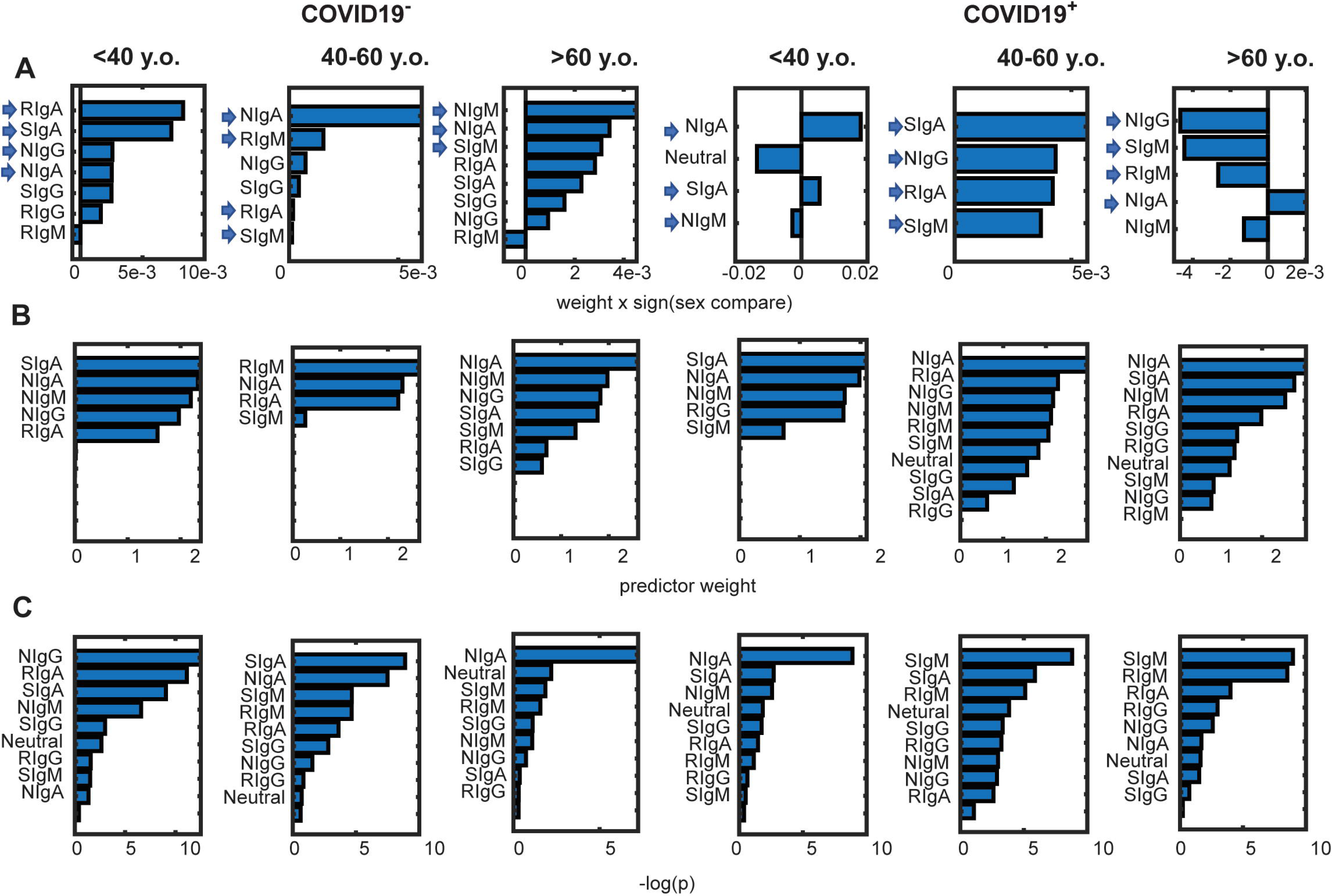
Selection of antibodies showing the largest difference between male and female cohorts in different age and acquired immunity subgroups. Three different feature selection methods were used to identify sex-differences in antibody responses in each of the six sample subgroups separated based on the COVID19 status and participants’ age (<40, 40-60, and >60 y.o.). Results from the **A)** Relieff machine learning feature selection method, **B)** feature selection method using neighborhood component analysis for regression (*fsrnca*), and **C)** univariate feature ranking for regression using F-tests (*fsrftest*) method, show major antibody measures that distinguish males from females within specified subgroups. Solid arrows indicate features selected by all three methods. Relieff plots show only antibody features with positive weight (i.e. features that contribute to significant sex separation) and the sign (i.e., +/-) in this representation is related to the male to female mean group ratios with positive sign indicating larger mean value in a male group.

It is important to observe that SIgM and NIgA come up as significant in 4 age and immunity subgroups and both of these antibodies show different clustering patterns in males and females (Figure 2). In addition to overall differences in these antibody features established by the three methods of feature selection analysis (Figure 3), there are significant changes in antibody responses over this time period. In males, there is a rise in the RIgA, NIgA, and SIgA serum levels generally following dose 2 vaccination. In contrast, higher IgM serum levels are noted in females over 60-year-old in both vaccination 1 and vaccination 2 periods.

### Correlation analysis provides quantitative links between sex and antibody responses

Further investigation of the relationship between serological profile and sex was performed using point-biserial correlation specifically developed for analysis of correlations between binary and continuous variables. Figure 5 shows antibody responses correlate with subjects’ sex. Once again, SIgM and RIgM are positively correlated with sex in individuals over 60 years old following both dose 1 and dose 2 vaccination. As is typical of antibody response evolution through Ig isotype switching with repeated vaccination (or even repeated infection), IgM levels show a negative correlation with sex in the younger age groups with values higher in males than in females.

**Figure 4.**
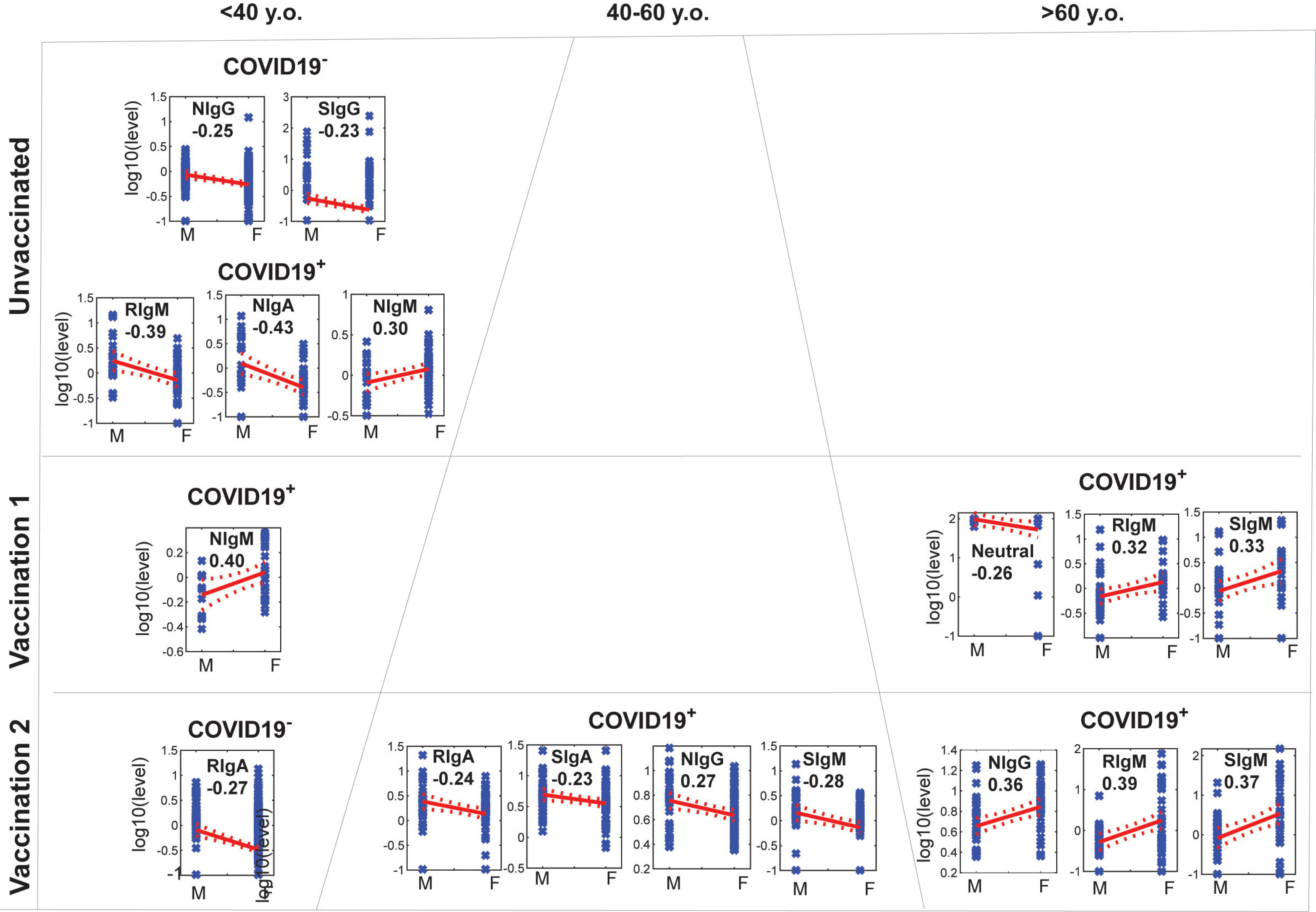
Direct correlation between antibody level and subjects’ sex. Point-biserial correlation between sex and serological factors. Shown are correlations with p<0.05 and absolute correlation value >0.20 within subgroups. A correlation value between sexes and log10 transformed concentration levels is indicated.

**Figure 5.**
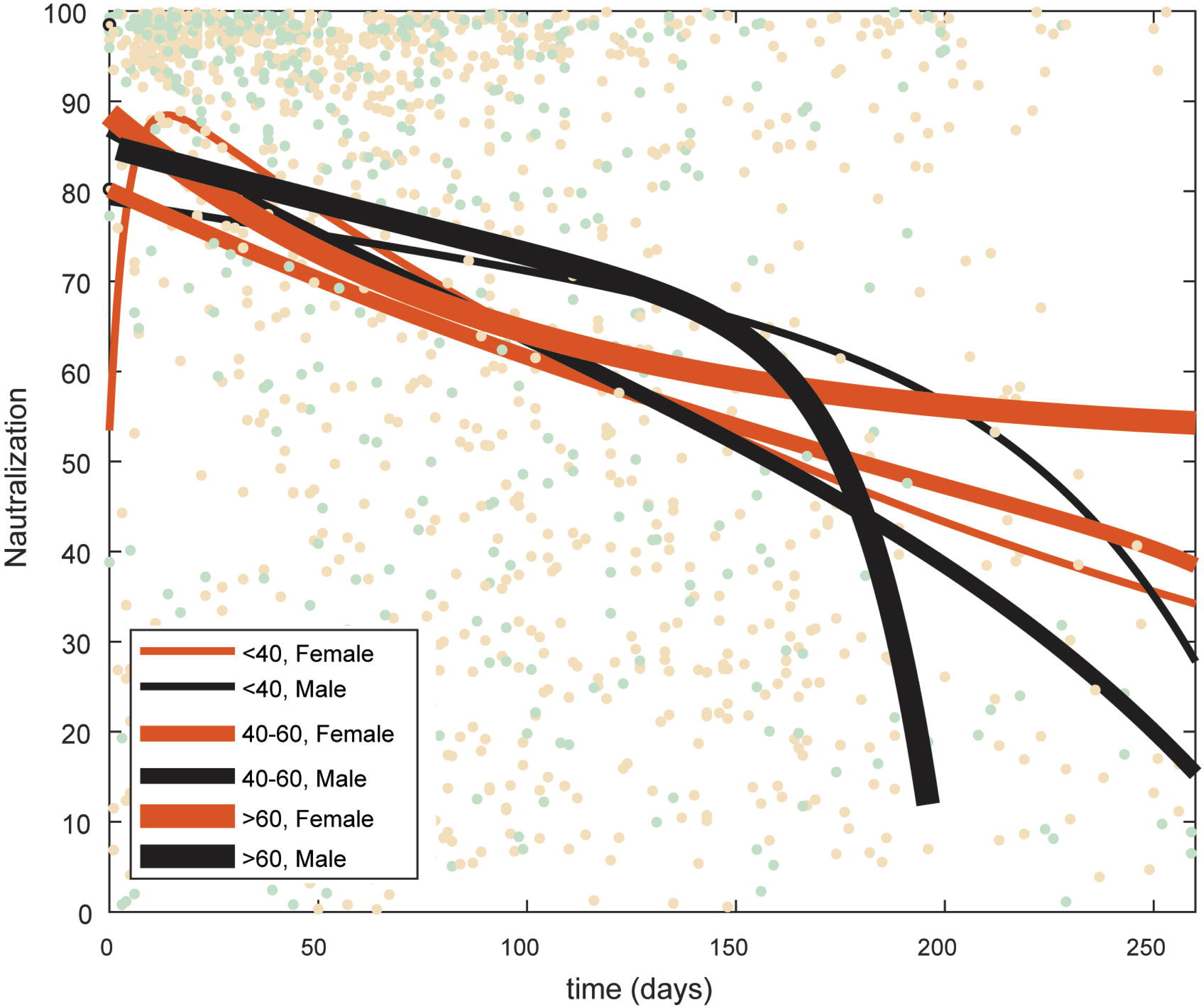
Antibody neutralization levels for over 60-year-old males decays faster than for other sex and age groups. Values present neutralization measurements in males and females in 3 different age groups following the second dose of COVID19 vaccination. Exponential fit is optimized separately for each represented group. Neutralization following Vaccination 2 for males and females. Shown are exponential best fit lines (as a*exp(b*time)+c*exp(d*time)). Time is shown in days from Vaccine 2. Y-axis shows the original value. Lines represent dark green, male, and red, female. Also included are actual sample measures showing major individual differences.

In the point-biserial correlation analysis, antibody neutralization, an important indicator of protection against serious COVID19 disease outcomes^31,32^ is only statistically significantly correlated with sex in the >60 group (Figure 4).

However, time course analysis (Figure 5) as well as feature selection (Figure 3) indicate some differences in neutralization trends between males and females. To determine whether there are specificities in the contributions of antibodies to neutralization in age and sex groups, a distance correlation analysis was performed between antibody neutralization measurements and other antibody response features as a measure of co-behaviour. In addition, relative square distance correlation was tested as a measure of contribution of each antibody response factor to the neutralization value. This analysis showed that after Vaccination 2, neutralization potential decays fastest among older persons who are male than any other group. This co-behaviour also suggests that females retain antibody-neutralization correlations longer than their male counterparts in all age groups.

### Correlation between antibodies and neutralization varies over time differently in males and females

Square of the correlation between two features represents a measure of the contribution of one feature to the variance of the other. Analysis of distance correlation between neutralization and each antibody in male and female groups over time following vaccination dose 1 and dose 2 shows major differences in the pattern of antibody contribution to neutralization (Figure 6). As an example, the contribution of RIgG and NIgA to neutralization in males and females under 40 years old are positively correlated over time and highly comparable by sex. However, neutralization and SIgM and RIgM are negatively correlated over time in these same groups. In contrast, following Dose 2 vaccination in those over 60 y.o. neutralization is similarly related to RIgG in males and females but inversely correlated with SIgG in the two groups. Thus, although the overall pattern in neutralization measurements is not correlated with sex, the observed levels of neutralization show different and complex relationships with antibody responses. Furthermore, analysis shown in Figure 6 suggests changes in the correlation of individual antibody responses over time, differing between males and females and by age group. In the under 40-year-old male group following vaccination 1, neutralization is initially strongly correlated to RIgG and NIgA concentrations and then more strongly to SIgG, NIgG, IgA and IgM antibodies. In females of this age group assessed at earlier time points, the relationship with neutralization is strongest with SIgG, NIgG and SIgA and all three IgM specificities. Following administration of Vaccination 2 in this group, the contribution of RIgG and SIgG to neutralization initially, at early time points post vaccination, drops.

**Figure 6.**
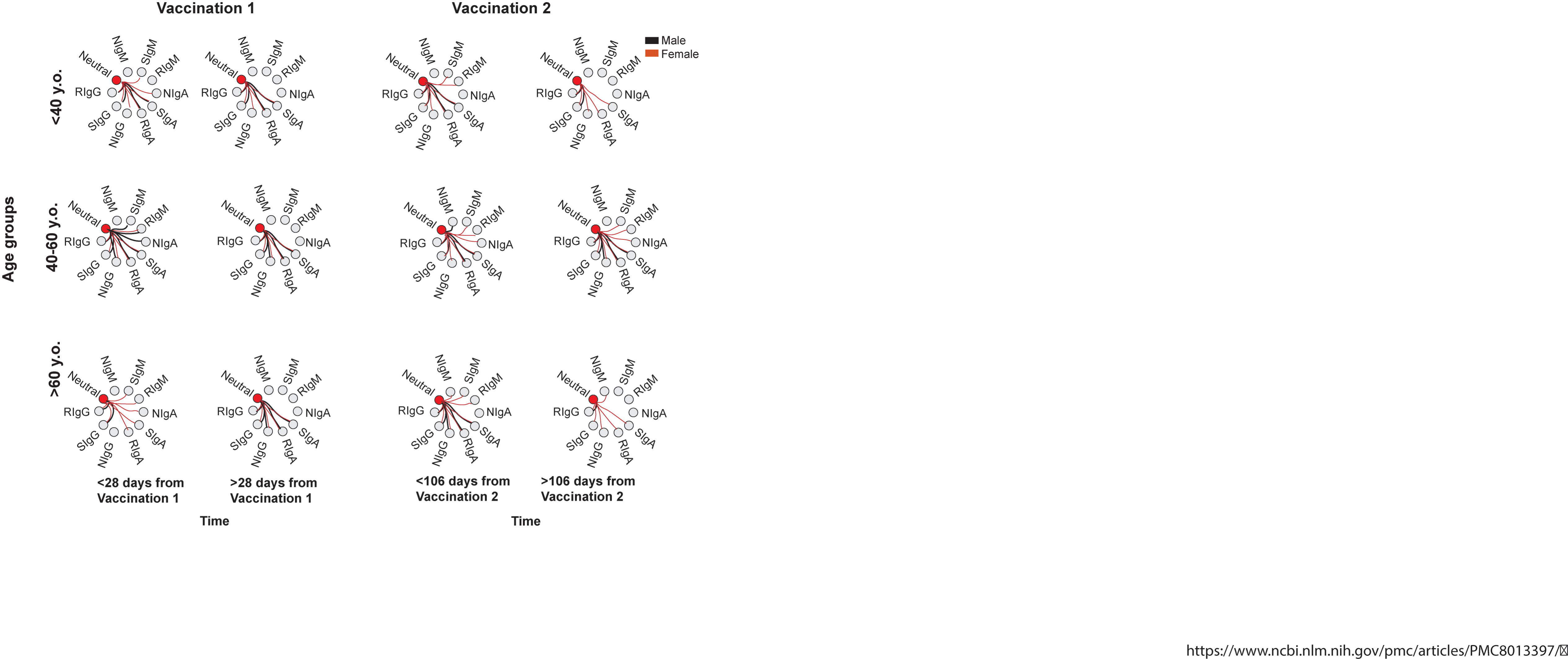
Correlation between neutralization and antibody isotypes is different in males and females in distinct age groups. Distance correlation between neutralization and all other features (pairwise) is calculated and thresholds were set using sample size appropriate correlation thresholds and p-values below 0.003 for measurements grouped by sex and age and time from vaccination. Vaccination groups include samples collected prior to half time between dose 1 and 2 (Vaccination 1) and dose 2 and 3 (Vaccination 2). Shown are correlation edges that pass the hard threshold separately for males (black line) and females (dark red line).

Changes in the contribution of antibody response features to neutralization over time in different groups can be further understood by exploring correlations in different time periods. For ease of presentation and increased number of samples in this case, time was binned into 21-day periods. Distance correlation was calculated using samples for each period and the square of individual values is divided with the sum of values for each time bin to provide a relative contribution. Thus, Figure 7 shows relative square distance correlation contributions to neutralization of each of the nine antibody response factors at 4 time points for each period. This representation is useful in that it highlights sex and age differences that were otherwise less apparent in an unsupervised PCA (Figure 7).

**Figure 7.**
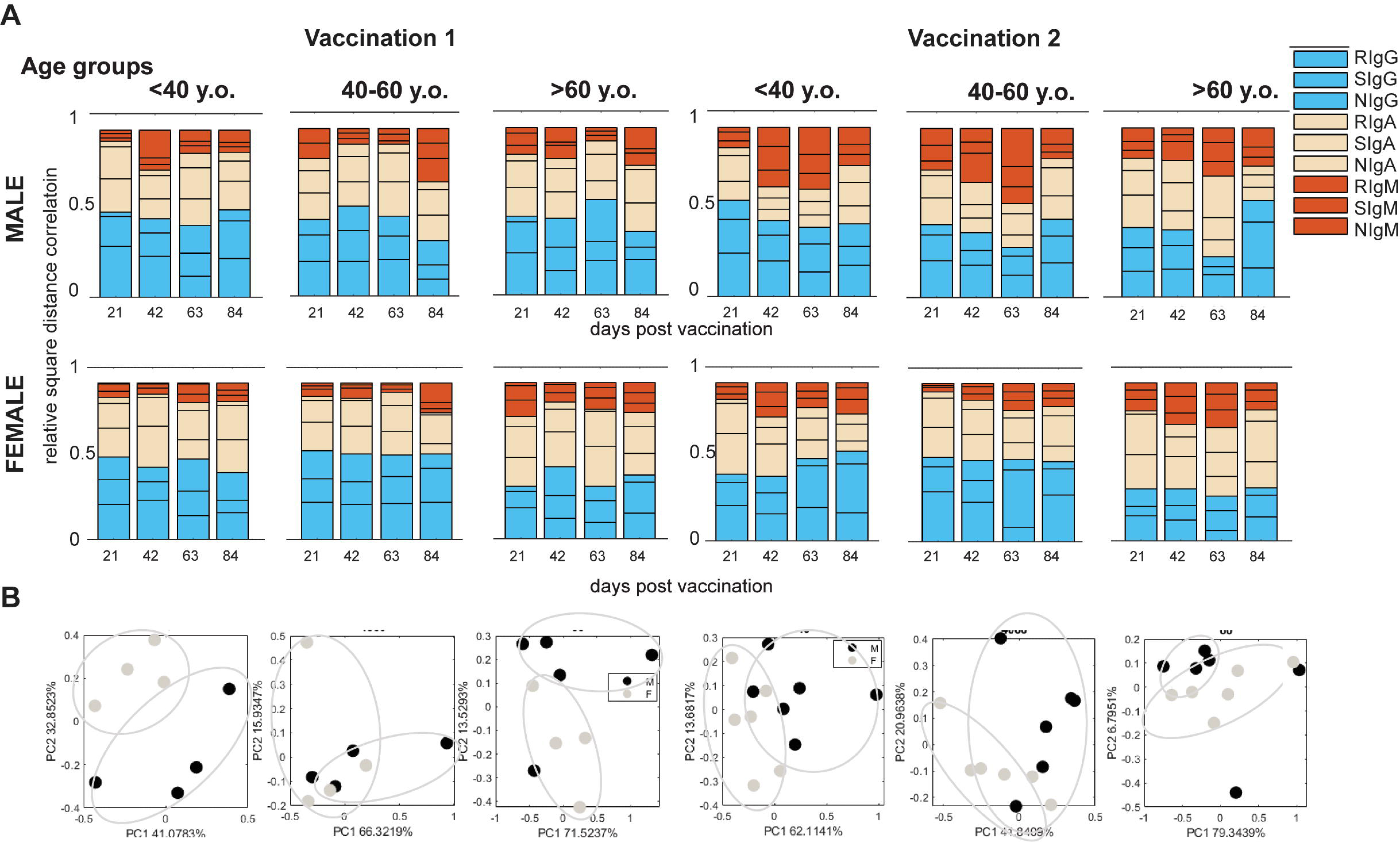
All measured antibody responses contribute to neutralization values at a level that is dependent on sex and age and changes over time. Relative square distance correlation (Y-axis) between each antibody response measured and neutralization was calculated for 21-day periods post Vaccination 1 and post Vaccination 2 in each subgroup of patients. COVID19 positive and negative subjects are combined in this analysis. Data are shown in bar graphs with relative contributions by immunoglobulin isotype responses (colour-coded) over time (days 21, 42, 63, and 84 post-Vaccination 2), and sex effects are depicted in plot groupings. Unsupervised PCAs show separation of males and females square correlation profiles at 4 or 5, 21-day periods in vaccination 1 and 2 (as indicated) for different age groups. Ellipsoids indicate visual groupings of male and female data points.

This contribution of different antibodies to neutralization over time depends on age and sex with differences observed in immunoglobulin isotype responses to all specific targeted proteins measured. With this novel approach to data analysis, we show that the relationship between antibody responses with neutralization depends on the subject’s age and sex, even in cases where the actual value of neutralization is not significantly different between these subgroups. As an example, following Vaccination 2 in the oldest group, the contribution of IgG is smaller than in younger groups particularly in females, while in under 40-year-olds males, contribution of IgMs is more significant than in females.

## Discussion

We applied novel network analyses to a complex set of antibody response data collected longitudinally from a large cohort of individuals over an important period of the COVID19 pandemic, bridging naïve immunity, infection-acquired immunity and immunity afforded across serial vaccination. The size and breadth of the cohort age range offered robust analytical outcomes by sex and age across multiple antibody parameters and repeated measures. Sex differences emerged with t-SNE analysis demonstrating that complex, non-linear analyses of these data would be required to understand factors driving antibody responses to COVID19. Correlations revealed important sex-associated co-behaviours of antibody responses with neutralization, some of which suggests young females may have stronger linkages in mucosal immunity and protection as their serum IgA responses are associated well with neutralization levels. Feature selection methods continued to confirm clustering analysis findings by age, with particularly prominent sex effects in IgA and NIgG responses in individuals <60. The capacity for significant Ig isotype switching with repeated vaccination was found to be weaker in older females whereas IgA responses in males could improve after Vaccination 2. This phenomenon was extended to an appreciation for the rate of antibody decay, which is quite rapid in males >60 and where females exhibited a superior retention of Ig isotype switching correlating with neutralization over time. Complex patterns of antibody response profiles in relation to neutralization differed by sex and age, as shown by distance correlations and these changed over time. These analytical approaches afforded several different angles by which to evaluate the complex relationships between antibody responses and neutralization showing that considerations for sex and age were highly important.

Demographic factors including age and sex are known to be associated with responses to both viral infection and immunization^1,20,33^. Our data shows differences in antibody responses across vaccination periods, in convalescent individuals with SARS-CoV-2 acquired immunity compared to individuals that did not show naturally acquired SARS-CoV2 immunity. Understanding the extent and type of this association and its change over time in relation to the resilience to an infection is crucial in attempts towards achieving more protective immunization and effective antiviral approaches. Extensive work on COVID19 immune responses led to an appreciation that antigen-specificity and Ig isotype can contribute to ACE2 neutralization^11,34,35^. While typical primary immune responses dominated by IgM evolve into recall immune responses composed of IgG, IgA, and IgE antibodies, there is variation in which factors contribute to protection against SARS-CoV-2, predictable in part based on antibody neutralization levels. Serum IgA is a dominant antibody soon after the onset of symptoms which peaks during the third week of the disease and provides the greatest contribution to neutralization. Assessment of bronchial alveolar fluid suggests this contributes predominantly to antibody-mediated virus neutralization activity in the lung compared to IgG^36^. Additionally, Ruggiero et al. showed that a coordinated response implicating both SIgM and SIgG is associated with protective immunity following vaccination^35^. While sex differences have been reported in disease outcomes, whether complex features of antibody responses to infection and vaccination play a role in determining immune protection to SARS-CoV-2 remains to be determined.

To identify potential differences in the coordination of the response over time in males and females of different ages, we have developed a novel methodology for analysis of correlation over time and applied it to a large longitudinal dataset spanning the COVID19 pandemic period, prior to and through vaccination dose series, with robust representation of subjects across different age groups and in both sexes. Our regression analysis of the age dependence of the measured serology profile is consistent with earlier studies showing age of ∼40 and ∼60 as significant points in the relationship between serology and response to the virus. Analyses of major features separating measurements in males and females in different age groups (<40 y.o, 40-60 y.o. and >60 y.o.) show changes at different points in the vaccination series, thus indicating differences in progression of antibody responses over time. In males, RIgA and SIgA show significantly faster increases following vaccination while the opposite is observed in SIgM and RIgM, particularly in the <40 y.o. and >60 y.o. groups. Although many selected antibodies have low weight in feature selection analysis, some are significantly correlated with subject sex. This is demonstrated in point-biserial correlation for different age and vaccination dose groups. RIgM is significantly correlated with sex in both young adult and senior groups but with opposite ratios, where RIgM is higher in males <r40 y.o. yet lower in females >60 y.o. Similarly, SIgM and NIgG are significantly correlated with sex in 40-60 years old and over 60 years old but are higher in males in the 40–60 y.o.- and in females in the over >60 y.o. group. However, neutralization levels are not correlated with sex in any of the age or vaccination groups suggesting different contributors to neutralization. To determine the relationship between neutralization and antibody production in these groups, we first performed a cluster analysis which revealed sex differences in the grouping of antibodies and neutralization in different groups. Furthermore, distance correlation and squared distance correlation analyses across 21-day period bins were employed to determine the contribution of different antibodies to neutralization. The level of correlation contribution was found to differ for different antibody responses over time and by sex and age. Squared distance correlation, representing the relative contribution of a variable to the level of another variable, was also calculated for each antibody measurement pairwise to normalization. Representation of this analysis shows change of contributing factors levels over time. Importantly, contributions in males and females in different age groups are sufficiently distinct allowing PCA separation of male and female samples based on this information. Unlike in the direct analysis of antibody serum where the data based on sex were not clearly separated by PCA, analysis of contributions of these factors to neutralization is clearly very different between sexes.

In sum, our study exploits novel, multivariate, machine learning, and network analysis tools integrating complex immunological parameters over time and across a diverse population to delve into the temporal dynamics of antibody responses through the COVID19 pandemic. Specifically, unsupervised and supervised machine learning strategies, coupled with square distance correlation, have enabled us to describe changes in the role of individual antibody responses in neutralization of ACE2. Contribution program changes over time and vaccination dose clearly differs between sex and age groups with major differences in infected vs. uninfected cohorts. In addition to the anticipated effects of age on the retention antibody responses, our analyses suggest that female antibody responses to SARS-CoV-2 vaccination may be of superior quality in their co-behaviour with protection (i.e., neutralization). Our work demonstrates that understanding antibody response profiles and their role in predicting neutralization goes beyond merely measuring absolute titres. Thus, rather than aiming for antibody titer achievements and changing only individual factors therein, our analyses supports the pursuit of optimized vaccination strategies that aim to alter the complete network of antibody responses. This sophisticated analysis of antibody responses also provides guidance for the parallel study of cell-mediated immunity where sex and age considerations are accounted for. There is an urgent need for such highly integrated indices of antibody and cell-mediated immunity as the search for reliable correlates of protection against COVID19 and other emerging pathogens continues, to best inform the design of highly effective vaccine approaches, and aid in clinical care.

## Conclusion

Using a variety of novel tools for knowledge discovery in datasets, we show that age and sex are important variables which impact the nature and temporal dynamics of infection-induced, vaccination-induced and hybrid immunity. Overall, this work describes a novel analytical pipeline with the potential to pinpoint changes in the response to infection and/or vaccination in the general or immune-compromised individuals.

---

**Supplementary Figure 1.** A. Principal Component Analysis (PCA) for dataset of 9 different antibody response and relative neutralization measurements with samples separated by immunity status, sex and vaccination period: B. tSNE visualization of overall sample differences. Groups are separated by age and COVID19 status as well as vaccination period. C. Table shows p-values for t-test comparison of difference between Males and Females in each projection group. Indicated in green are statistically significant differences.

**Supplementary Figure 2.** Hierarchical clustering of subjects divided into groups by sex and COVID19 status using 9 different antibody response and relative neutralization measurements. A major dependence on vaccination period is indicated (legend indicates vaccination state: Unvaccinated prior to vaccination, Vaccination 1 after first dose and prior to dose 2 etc.). Shown is a relative value heath map with blue showing lower and red showing and higher than mean values.

## References

1. Scully, E.P., Haverfield, J., Ursin, R.L., Tannenbaum, C., and Klein, S.L. (2020). Considering how biological sex impacts immune responses and COVID-19 outcomes. Nat Rev Immunol 20. 10.1038/s41577-020-0348-8.

2. Arnold, C.G., Libby, A., Vest, A., Hopkinson, A., and Monte, A.A. (2022). Immune mechanisms associated with sex-based differences in severe COVID-19 clinical outcomes. Preprint, 10.1186/s13293-022-00417-3.

3. Takahashi, T., Ellingson, M.K., Wong, P., Israelow, B., Lucas, C., Klein, J., Silva, J., Mao, T., Oh, J.E., Tokuyama, M., et al. (2020). Sex differences in immune responses that underlie COVID-19 disease outcomes. Nature 588. 10.1038/s41586-020-2700-3.

4. Dehingia, N., and Raj, A. (2021). Sex differences in COVID-19 case fatality: do we know enough? Preprint, 10.1016/S2214-109X(20)30464-2.

5. 5. Nguyen, N.T., Chinn, J., de Ferrante, M., Kirby, K.A., Hohmann, S.F., and Amin, A. (2021). Male gender is a predictor of higher mortality in hospitalized adults with COVID-19. PLoS One 16. 10.1371/journal.pone.0254066.

6. Alghamdi, I.G., Hussain, I.I., Almalki, S.S., Alghamdi, M.S., Alghamdi, M.M., and El-Sheemy, M.A. (2014). The pattern of Middle east respiratory syndrome coronavirus in Saudi Arabia: A descriptive epidemiological analysis of data from the Saudi Ministry of Health. Int J Gen Med 7. 10.2147/IJGM.S67061.

7. Channappanavar, R., Fett, C., Mack, M., Ten Eyck, P.P., Meyerholz, D.K., and Perlman, S. (2017). Sex-Based Differences in Susceptibility to Severe Acute Respiratory Syndrome Coronavirus Infection. The Journal of Immunology 198. 10.4049/jimmunol.1601896.

8. Peckham, H., de Gruijter, N.M., Raine, C., Radziszewska, A., Ciurtin, C., Wedderburn, L.R., Rosser, E.C., Webb, K., and Deakin, C.T. (2020). Male sex identified by global COVID-19 meta-analysis as a risk factor for death and ITU admission. Nat Commun 11. 10.1038/s41467-020-19741-6.

9. Lott, N., Gebhard, C.E., Bengs, S., Haider, A., Kuster, G.M., Regitz-Zagrosek, V., and Gebhard, C. (2023). Sex hormones in SARS-CoV-2 susceptibility: key players or confounders? Preprint, 10.1038/s41574-022-00780-6.

10. Taneja, V. (2018). Sex hormones determine immune response. Front Immunol 9. 10.3389/fimmu.2018.01931.

11. Klingler, Jã., Weiss, S., Itri, V., Liu, X., Oguntuyo, K.Y., Stevens, C., Ikegame, S., Hung, C.T., Enyindah-Asonye, G., Amanat, F., et al. (2021). Role of Immunoglobulin M and A Antibodies in the Neutralization of Severe Acute Respiratory Syndrome Coronavirus 2. Journal of Infectious Diseases 223. 10.1093/infdis/jiaa784.

12. 12. Goodhue Meyer, E., Simmons, G., Grebe, E., Gannett, M., Franz, S., Darst, O., Di Germanio, C., Stone, M., Contestable, P., Prichard, A., et al. (2021). Selecting COVID-19 convalescent plasma for neutralizing antibody potency using a high-capacity SARS-CoV-2 antibody assay. Transfusion (Paris) 61. 10.1111/trf.16321.

13. Diao, B., Wang, C., Tan, Y., Chen, X., Liu, Y., Ning, L., Chen, L., Li, M., Liu, Y., Wang, G., et al. (2020). Reduction and Functional Exhaustion of T Cells in Patients With Coronavirus Disease 2019 (COVID-19). Front Immunol 11. 10.3389/fimmu.2020.00827.

14. Giamarellos-Bourboulis, E.J., Netea, M.G., Rovina, N., Akinosoglou, K., Antoniadou, A., Antonakos, N., Damoraki, G., Gkavogianni, T., Adami, M.E., Katsaounou, P., et al. (2020). Complex Immune Dysregulation in COVID-19 Patients with Severe Respiratory Failure. Cell Host Microbe 27. 10.1016/j.chom.2020.04.009.

15. Grifoni, A., Weiskopf, D., Ramirez, S.I., Mateus, J., Dan, J.M., Moderbacher, C.R., Rawlings, S.A., Sutherland, A., Premkumar, L., Jadi, R.S., et al. (2020). Targets of T Cell Responses to SARS-CoV-2 Coronavirus in Humans with COVID-19 Disease and Unexposed Individuals. Cell 181. 10.1016/j.cell.2020.05.015.

16. Peng, Y., Mentzer, A.J., Liu, G., Yao, X., Yin, Z., Dong, D., Dejnirattisai, W., Rostron, T., Supasa, P., Liu, C., et al. (2020). Broad and strong memory CD4+ and CD8+ T cells induced by SARS-CoV-2 in UK convalescent individuals following COVID-19. Nat Immunol 21. 10.1038/s41590-020-0782-6.

17. Del Valle, D.M., Kim-Schulze, S., Huang, H.H., Beckmann, N.D., Nirenberg, S., Wang, B., Lavin, Y., Swartz, T.H., Madduri, D., Stock, A., et al. (2020). An inflammatory cytokine signature predicts COVID-19 severity and survival. Nat Med 26. 10.1038/s41591-020-1051-9.

18. Lucas, C., Wong, P., Klein, J., Castro, T.B.R., Silva, J., Sundaram, M., Ellingson, M.K., Mao, T., Oh, J.E., Israelow, B., et al. (2020). Longitudinal analyses reveal immunological misfiring in severe COVID-19. Nature 584. 10.1038/s41586-020-2588-y.

19. Leung, G.M., Medley, A.J., Ho, L.M., Chau, P., Wong, I.O.L., Thach, T.Q., Ghani, A.C., Donnelly, C.A., Fraser, C., Riley, S., et al. (2004). The epidemiology of severe acute respiratory syndrome in the 2003 Hong Kong epidemic: An analysis of all 1755 patients. Ann Intern Med 141. 10.7326/0003-4819-141-9-200411020-00006.

20. Fink, A.L., Engle, K., Ursin, R.L., Tang, W.Y., and Klein, S.L. (2018). Biological sex affects vaccine efficacy and protection against influenza in mice. Proc Natl Acad Sci U S A 115. 10.1073/pnas.1805268115.

21. Khan, S.R., van der Burgh, A.C., Peeters, R.P., van Hagen, P.M., Dalm, V.A.S.H., and Chaker, L. (2021). Determinants of Serum Immunoglobulin Levels: A Systematic Review and Meta-Analysis. Front Immunol 12. 10.3389/fimmu.2021.664526.

22. Khan, S.R., Chaker, L., Ikram, M.A., Peeters, R.P., van Hagen, P.M., and Dalm, V.A.S.H. (2021). Determinants and Reference Ranges of Serum Immunoglobulins in Middle-Aged and Elderly Individuals: a Population-Based Study. J Clin Immunol 41. 10.1007/s10875-021-01120-5.

23. Sha, C., Cuperlovic-Culf, M., and Hu, T. (2021). SMILE: systems metabolomics using interpretable learning and evolution. BMC Bioinformatics 22. 10.1186/s12859-021-04209-1.

24. Cuperlovic-Culf, M., Nguyen-Tran, T., and Bennett, S.A.L. (2023). Machine Learning and Hybrid Methods for Metabolic Pathway Modeling. In Methods in Molecular Biology 10.1007/978-1-0716-2617-7_18.

25. Monti, F., Stewart, D., Surendra, A., Alecu, I., Nguyen-Tran, T., Bennett, S.A.L., and Cuperlovic-Culf, M. (2023). Signed Distance Correlation (SiDCo): an online implementation of distance correlation and partial distance correlation for data-driven network analysis. Bioinformatics 39. 10.1093/bioinformatics/btad210.

26. Collins, E., Galipeau, Y., Arnold, C., Bosveld, C., Heiskanen, A., Keeshan, A., Nakka, K., Shir-Mohammadi, K., St-Denis-Bissonnette, F., Tamblyn, L., et al. (2022). Cohort profile: S top the Spread Ottawa (SSO) - a community-based prospective cohort study on antibody responses, antibody neutralisation efficiency and cellular immunity to SARS-CoV-2 infection and vaccination. BMJ Open 12. 10.1136/bmjopen-2022-062187.

27. Colwill, K., Galipeau, Y., Stuible, M., Gervais, C., Arnold, C., Rathod, B., Abe, K.T., Wang, J.H., Pasculescu, A., Maltseva, M., et al. (2022). A scalable serology solution for profiling humoral immune responses to SARS-CoV-2 infection and vaccination. Clin Transl Immunology 11. 10.1002/cti2.1380.

28. Belacel, N., Čuperlović-Culf, M., Laflamme, M., and Ouellette, R. (2004). Fuzzy J-Means and VNS methods for clustering genes from microarray data. Bioinformatics 20. 10.1093/bioinformatics/bth142.

29. Reese, S.E., Archer, K.J., Therneau, T.M., Atkinson, E.J., Vachon, C.M., De Andrade, M., Kocher, J.P.A., and Eckel-Passow, J.E. (2013). A new statistic for identifying batch effects in high-throughput genomic data that uses guided principal component analysis. Bioinformatics 29. 10.1093/bioinformatics/btt480.

30. Székely, G.J., Rizzo, M.L., and Bakirov, N.K. (2007). Measuring and testing dependence by correlation of distances. Ann Stat 35. 10.1214/009053607000000505.

31. Ju, B., Zhang, Q., Ge, J., Wang, R., Sun, J., Ge, X., Yu, J., Shan, S., Zhou, B., Song, S., et al. (2020). Human neutralizing antibodies elicited by SARS-CoV-2 infection. Nature 584. 10.1038/s41586-020-2380-z.

32. Piccoli, L., Park, Y.J., Tortorici, M.A., Czudnochowski, N., Walls, A.C., Beltramello, M., Silacci-Fregni, C., Pinto, D., Rosen, L.E., Bowen, J.E., et al. (2020). Mapping Neutralizing and Immunodominant Sites on the SARS-CoV-2 Spike Receptor-Binding Domain by Structure-Guided High-Resolution Serology. Cell 183. 10.1016/j.cell.2020.09.037.

33. Klein, S.L., and Flanagan, K.L. (2016). Sex differences in immune responses. Preprint, 10.1038/nri.2016.90.

34. Ma, H., Zeng, W., He, H., Zhao, D., Jiang, D., Zhou, P., Cheng, L., Li, Y., Ma, X., and Jin, T. (2020). Serum IgA, IgM, and IgG responses in COVID-19. Preprint, 10.1038/s41423-020-0474-z.

35. Ruggiero, A., Piubelli, C., Calciano, L., Accordini, S., Valenti, M.T., Carbonare, L.D., Siracusano, G., Temperton, N., Tiberti, N., Longoni, S.S., et al. (2022). SARS-CoV-2 vaccination elicits unconventional IgM specific responses in naïve and previously COVID-19-infected individuals. EBioMedicine 77. 10.1016/j.ebiom.2022.103888.

36. Sterlin, D., Mathian, A., Miyara, M., Mohr, A., Anna, F., Claër, L., Quentric, P., Fadlallah, J., Devilliers, H., Ghillani, P., et al. (2021). IgA dominates the early neutralizing antibody response to SARS-CoV-2. Sci Transl Med 13. 10.1126/scitranslmed.abd2223.

