## Supplementary figures and images for "Multi-variate statistical and machine learning reveals the interplay between sex and age in antibody responses to *de novo* SARS-CoV-2 infection and vaccination"

### Supplementary_Figure 1..tiff

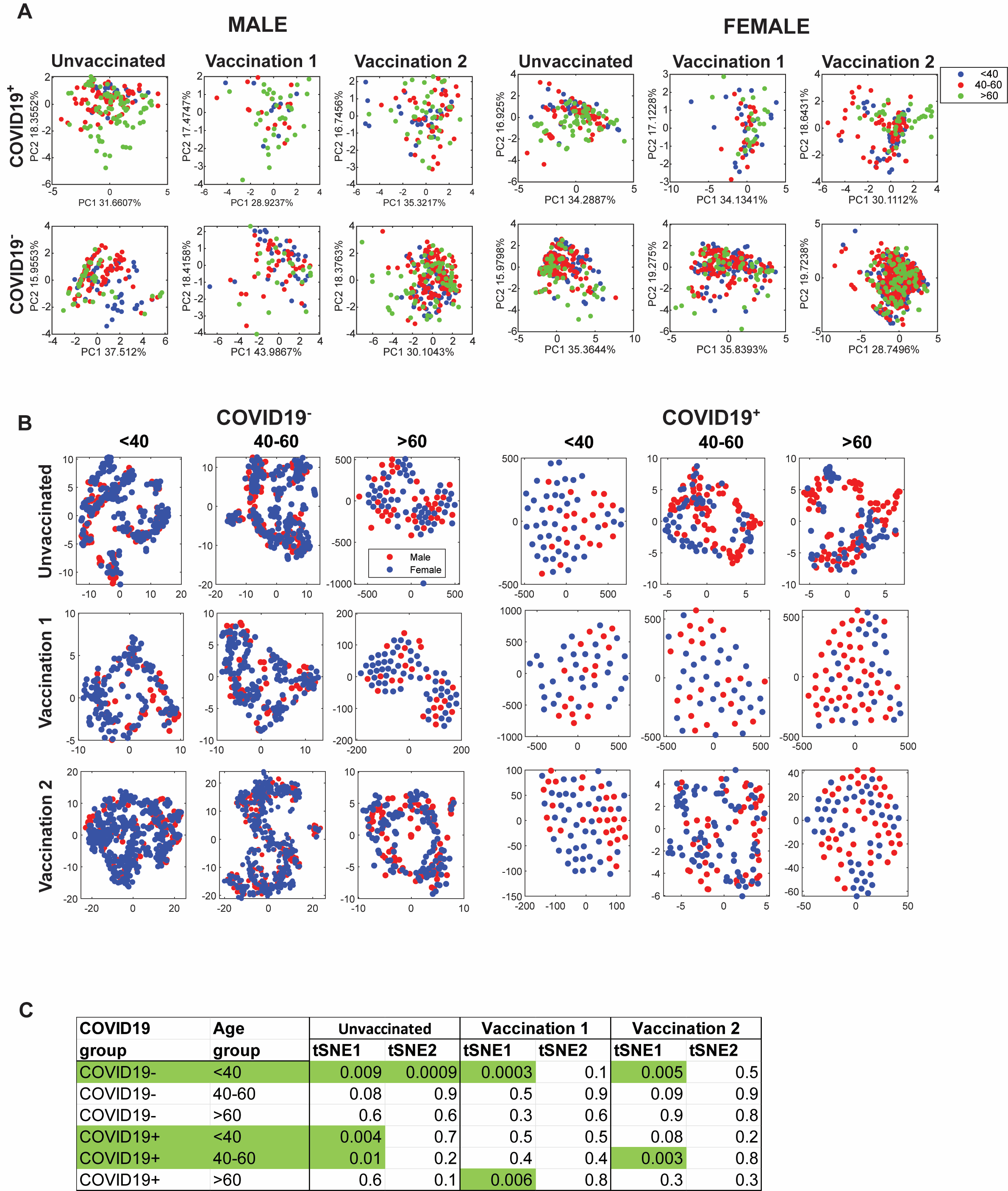

### Supplementary_Figure 2..tiff

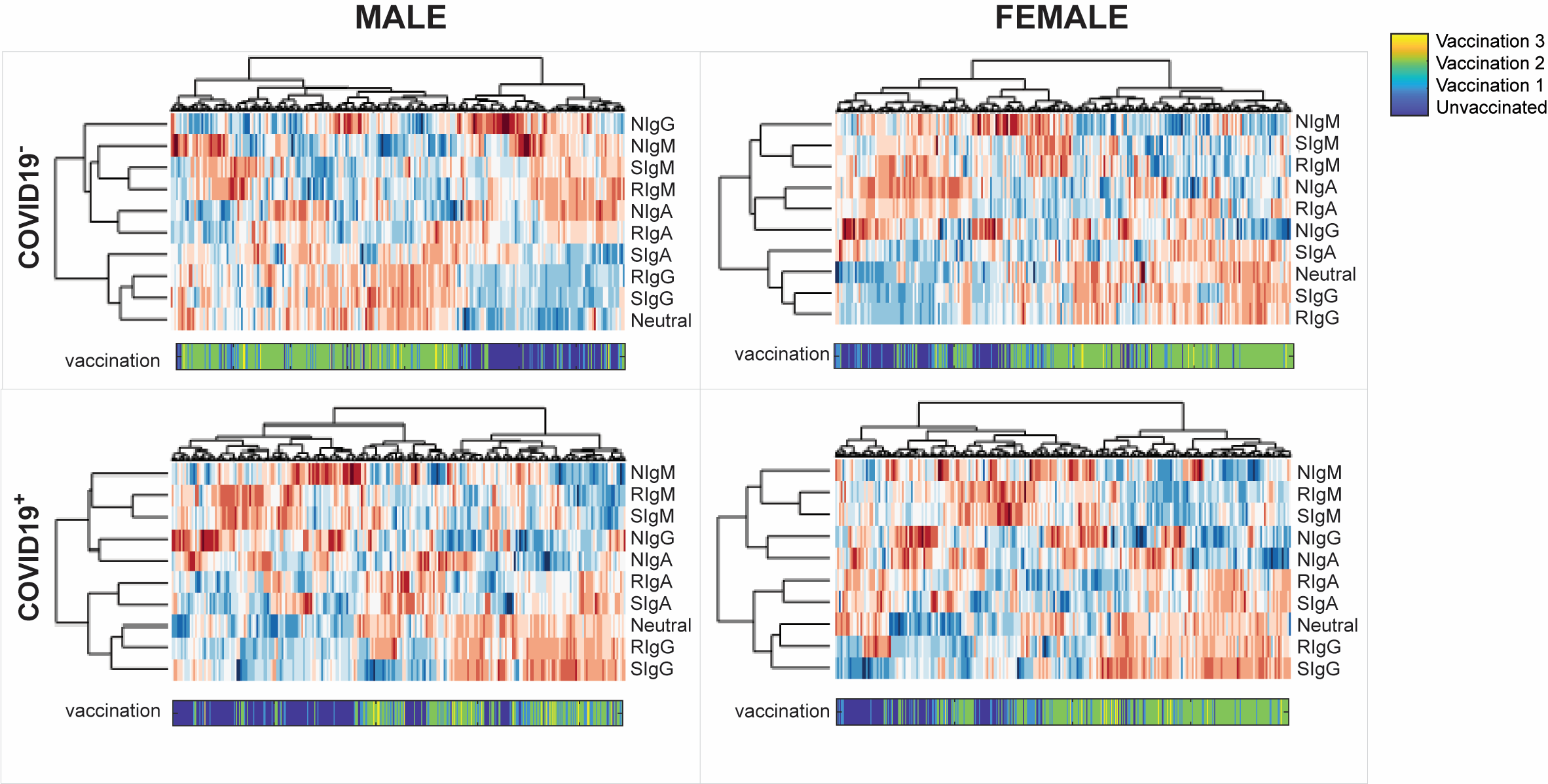
